# Recurrent Body Shape Patterns in Cave- and Surface-Dwelling Trichomycterid Catfishes

**DOI:** 10.64898/2026.03.03.709414

**Authors:** Nelson Falcón-Espitia, Carlos Daniel Cadena

## Abstract

The evolution of body shape reflects the interplay between functional constraints and habitat structure. While cave environments are well known to promote regressive traits in fish such as eye and pigment loss, their influence on overall body form remains poorly understood. Here, we examine patterns of body shape variation in cave-dwelling and surface-dwelling trichomycterid catfishes from northeastern Colombia to assess whether consistent associations exist between habitat type and morphology. Using geometric morphometric analyses, we quantified differences in body shape among species inhabiting subterranean and surface environments. Our results reveal significant habitat-associated differentiation in body shape associated with habitat along the main axes of morphological variation, despite some overlap indicating that habitat does not fully predict morphological variation. Cave-dwelling species exhibit more elongated and fusiform body shapes, whereas surface-dwelling species tend to have deeper and more robust morphologies. These patterns are consistent with differences in habitat-associated ecological conditions, although no direct functional or performance data were evaluated. The recurrence of similar body shapes among species from different clades occupying comparable habitats is consistent with repeated morphological responses to shared ecological constraints.

**Research Highlights:**

- Multivariate shape analyses reveal significant habitat-associated variation in trichomycterid fishes, with partial overlap indicating that habitat does not fully predict morphological variation.
- Body shape differs consistently between cave-and surface-dwelling trichomycterids: cave species exhibit more elongated, fusiform forms, whereas surface species display deeper body configurations.
- Recurrent body shape patterns across independent lineages are consistent with repeated phenotypic responses to shared ecological constraints, without directly testing functional or performance mechanisms.

## 1 Introduction

The evolutionary responses of organisms to environmental pressures arise from the interplay between natural selection, genetic architecture, and developmental and functional constraints (Arnold, 1992). Functional constraints can bias both the direction and magnitude of phenotypic change by mediating trade-offs among performance traits, thereby shaping how organisms respond to ecological gradients (Adams & Nistri, 2010; Black & Armbruster, 2022; Evans, Vidal-García, Tagliacollo, Taylor, & Fenolio, 2019; Walker, 2007). Rather than producing unlimited phenotypic outcomes, these constraints may channel variation along a restricted set of morphofunctional axes, generating recurrent associations between phenotype and environment (Grossnickle et al., 2024; Losos, 2011). Such recurrent associations are often interpreted as signatures of convergent evolution, even when similar phenotypes arise through different developmental or genetic pathways (Losos, 2011; Moen, 2019; Sansalone et al., 2020). In addition, empirical work in model systems such as threespine sticklebacks has shown that phenotypic variation along these axes may reflect locally adaptive responses to environmental gradients and, potentially, phenotypic plasticity, often involving integrated suites of traits rather than isolated morphological changes (Smith, Zięba, Spence, Klepaker, & Przybylski, 2020; Spence, Wootton, Barber, Przybylski, & Smith, 2013).

Aquatic locomotion provides a well-developed framework to explore these ideas, as body shape is tightly linked to hydrodynamic performance and habitat use. Classical models of fish locomotion predict a trade-off between sustained (steady) swimming efficiency and unsteady swimming performance, including acceleration and maneuverability (Langerhans, 2008; Langerhans & Reznick, 2010; Webb, 1984). Fusiform and elongate body shapes tend to reduce drag and improve efficiency during sustained swimming; traits typically associated with relatively open or hydrodynamically simple environments. In contrast, deeper and more laterally compressed bodies, often coupled with posteriorly positioned fins, enhance maneuverability and rapid acceleration, traits favored in structurally complex habitats where obstacle avoidance and fine-scale movement control are critical (Breda, Oliveira, & Goulart, 2005; Langerhans & Reznick, 2010; Webb, 1984). These expectations are grounded in the functional structure of habitats (such as flow variability, substrate heterogeneity, and presence of obstacles) rather than in broad habitat categories.

Subterranean ecosystems offer a unique opportunity to evaluate how such functional constraints may shape phenotypic evolution. Caves have been colonized by a wide diversity of organisms, including bacteria, fungi, invertebrates, and vertebrates, many of which display recurrent morphological, physiological, and behavioral traits associated with subterranean life (Lee et al., 2012; Martinelli-Marín, Lasso, Sauro, & Caballero-Gaitán, 2026; Soares & Niemiller, 2013, 2020; White & Culver, 2012). From a functional perspective, cave and surface aquatic environments may differ in key ecological dimensions such as flow regimes, spatial configuration, and patterns of movement and resource use, which can influence locomotor demands and the selective landscape experienced by organisms (Juan, Guzik, Jaume, & Cooper, 2010; Lasso, Barriga, & Fernández-Auderset, 2019). Research on cave-adapted organisms has traditionally emphasized so-called regressive traits (such as eye reduction and loss of pigmentation), often interpreted as responses to the absence of light and relaxed selection on visual systems (Juan et al., 2010; Mammola et al., 2020; Wilkens, 2010). In contrast, other dimensions of the phenotype that may reflect functional interactions with the physical environment including overall body shape and locomotor morphology, have received less attention in subterranean systems.

Cave-dwelling fishes exemplify this imbalance, with more than 300 species of ray-finned fish (Actinopterygii) belonging to multiple orders inhabiting subterranean systems worldwide (Soares & Niemiller, 2013). Among these species, the Neotropical Trichomycteridae family, comprising nearly 500 species distributed across a broad elevational and ecological range, is one of the most diverse to have repeatedly colonized cave environments (Fricke, Eschmeyer, & Fong, 2025; Martinelli-Marín et al., 2026). Several cave-dwelling species of the genus *Trichomycterus* exhibit pronounced troglomorphic traits, including eye reduction, depigmentation, and paedomorphic features (C. Castellanos-Morales, 2008; Flórez, Cadena, DoNascimiento, & Torres, 2021; Jeffery, 2001; Langecker & Longley, 1993; Soares & Niemiller, 2013, 2020; Trajano, 2021). Phylogenetic evidence further indicates that subterranean habitats have been colonized independently by multiple lineages within this group (Flórez et al., 2021; Trajano, 2021) providing an opportunity to examine whether similar habitat conditions are associated with comparable patterns of body shape variation.

Despite this growing body of work, it remains unclear whether adaptation to cave environments in trichomycterid fishes involves consistent modifications in body shape, beyond the classic regressive traits. Most studies on cave-dwelling *Trichomycterus* have relied on linear morphometrics for taxonomic purposes or qualitative descriptions of troglomorphism (e.g., C. Castellanos-Morales, 2007, 2008; Mesa, Lasso, Ochoa, & DoNascimiento, 2018), leaving unexplored how body shape varies in relation to habitat-specific functional demands. In other cavefish systems, such as *Astyanax* in Mexico and *Sinocyclocheilus* in China, studies have documented substantial and functionally relevant morphological divergence associated with subterranean life, including changes in cranial morphology, body depth, and overall body configuration linked to feeding strategies, sensory systems, and ecological specialization (Mao et al., 2021; Powers, Davis, Kaplan, & Gross, 2017; R. Reyes, 2022). Importantly, these systems also reveal that phenotypic responses to cave environments are not uniform, but instead encompass multiple morphological trajectories shaped by different ecological and evolutionary contexts. Accordingly, rather than providing direct tests of functional mechanisms, such studies offer a comparative framework to interpret patterns of morphological variation in relation to potential ecological drivers.

In the Eastern Andes of Colombia, extensive karst systems in the department of Santander host a diverse subterranean fauna, including multiple cave-dwelling species of *Trichomycterus* (A. Castellanos-Morales, Marino-Zamudio, Guerrero, & Maldonado-Ocampo, 2011; Lasso, Mesa, Castellanos-Morales, Fernández, & Do Nascimiento, 2018; Martinelli-Marín et al., 2026; Muñoz-Saba, González-Sánchez, & Calvo-Roa, 2013). Aquatic habitats within these cave systems are spatially confined and differ from surface streams in terms of resource availability, spatial configuration and, probably, patterns of organismal movement (Lasso et al., 2019). Rather than reflecting a single dimension of environmental variation, these differences likely involve multiple interacting factors, including hydrodynamics, habitat structure, and behavioral context, which together shape the functional demands experienced by fishes. However, the natural history of organisms living in these caves is very little known (Lasso et al., 2019).

In this study, we examine patterns of body shape variation among cave-dwelling and surface-dwelling trichomycterid catfishes from Colombia. Specifically, we assess whether body shape variation is consistently associated with habitat of occurrence (subterranean vs. surface) across species and lineages within a local geographical context. By comparing species inhabiting similar environments, we evaluate whether recurrent body configurations emerge in association with shared habitat conditions and interpret these patterns in the light of functional expectations related to habitat structure and locomotor demands. This approach allows us to explore whether habitat-associated constraints are linked to repeated phenotypic patterns across independent evolutionary lineages.

## 2 Materials and Methods

### Study system

*Trichomycterus* as currently defined is not a monophyletic genus, and it comprises a considerable number of species within Trichomycterinae classified into two major clades: *Trichomycterus* and *Eremophilus* (Fernandez, Arroyave, & Schaefer, 2021). Within this group, *Eremophilus mutisii*, a sighted and highly pigmented species inhabiting rivers and streams in the high Andean plateau known as the Altiplano Cundiboyacense, is the closest known relative of *Trichomycterus rosablanca*, a cave-dwelling species from the karstic system of Santander that exhibits pronounced troglomorphic traits characters (DoNascimiento & Prada-Pedreros, 2020; Flórez et al., 2021; Mesa et al., 2018). These two species represent contrasting ends of a troglomorphic gradient, with *T. rosablanca* exhibiting complete depigmentation and eye loss, whereas *E. mutisii* retains fully developed eyes and pigmentation.

In a different clade, *T. latistriatus* is most closely related to *T. sandovali*, forming a group in which pigmentation and eye development vary widely among individuals from cave and surface habitats (Flórez et al., 2021). Within this clade, *T. sandovali* typically exhibits reduced eye development and variable pigmentation associated with subterranean habits, whereas *T. latistriatus* shows greater development of both traits, although substantial variation exists among individuals.

Similarly, *T. ruitoquensis* exhibits intermediate characteristics, including variable pigmentation and reduced but functional eyes, and has been reported from both cave and surface environments. Together, these species illustrate a continuum of troglomorphic expression rather than discrete ecological categories. The morphological similarities of cave species in these groups suggest that adaptation to subterranean life may have arisen convergently in different lineages, leading to distinct phenotypic modifications in cave environments.

### Sampling

Our study examined multiple individuals representing cave and surface lineages of different *Trichomycterus* subclades (Flórez et al., 2021), all of which were sourced from biological collections and preserved following standard protocols (formalin fixation and subsequent storage in ethanol).

To enable comparisons within a local-scale ecological framework, specimens were grouped into habitat categories (cave and surface) based on their reported habitat of occurrence (Table 1). This classification was implemented prior to body shape analyses to explicitly test habitat-associated morphological variation. Although habitat assignment based on collection records may introduce some uncertainty, our categorization is consistent with current taxonomic and ecological knowledge of these species (Lasso et al., 2019, 2018).

**Table 1.** Composition of the dataset used for geometric morphometric analyses, showing the number of specimens examined per species and habitat. Phylogenetic group assignment follows Flórez et al. (2021). For species occurring in both cave and surface environments, sample sizes are reported separately for each habitat.

| Species | Phylogenetic group<br>(Flórez et al. 2021) | Habitat | N |
| --- | --- | --- | --- |
| *Trichomycterus rosablanca* | *rosablanca* | Cave | 70 |
| *Eremophilus mutisii* |  | Surface | 38 |
| *Trichomycterus sandovali* | *sandovali* | Cave | 46 |
| *Trichomycterus latistriatus* |  | Cave / Surface | 25 / 13 |
| *Trichomycterus ruitoquensis* | *ruitoquensis* | Cave / Surface | 7 / 37 |

Following DoNascimiento & Prada-Pedreros (2020), we considered *T. guacamayoensis* (all specimens from caves) to be synonymous with *T. latistriatus*. Additionally, for species with records in both habitat types (*T. latistriatus* and *T. ruitoquensis*), specimens were analyzed according to their specific habitat of origin (Table 1). The degree of connectivity among subterranean and surface populations in these systems remains uncertain; however, regional karst dynamics suggest the possibility of intermittent hydrological connectivity, at least seasonally (Lasso et al., 2019).

All sampling localities for species of the *Trichomycterus* genus are constrained to a single region, as cave systems and surface sites are located within the Santander department, northeastern Colombia (Fig 1). These correspond to small karstic drainages characterized by short surface streams and isolated hypogean systems (Flórez et al., 2021; Lasso et al., 2019; Muñoz-Saba et al., 2013). These systems exhibit structural and environmental heterogeneity, including both accessible passages and permanently submerged sections, and therefore should not considered uniformly aphotic environments.

**Figure 1.**
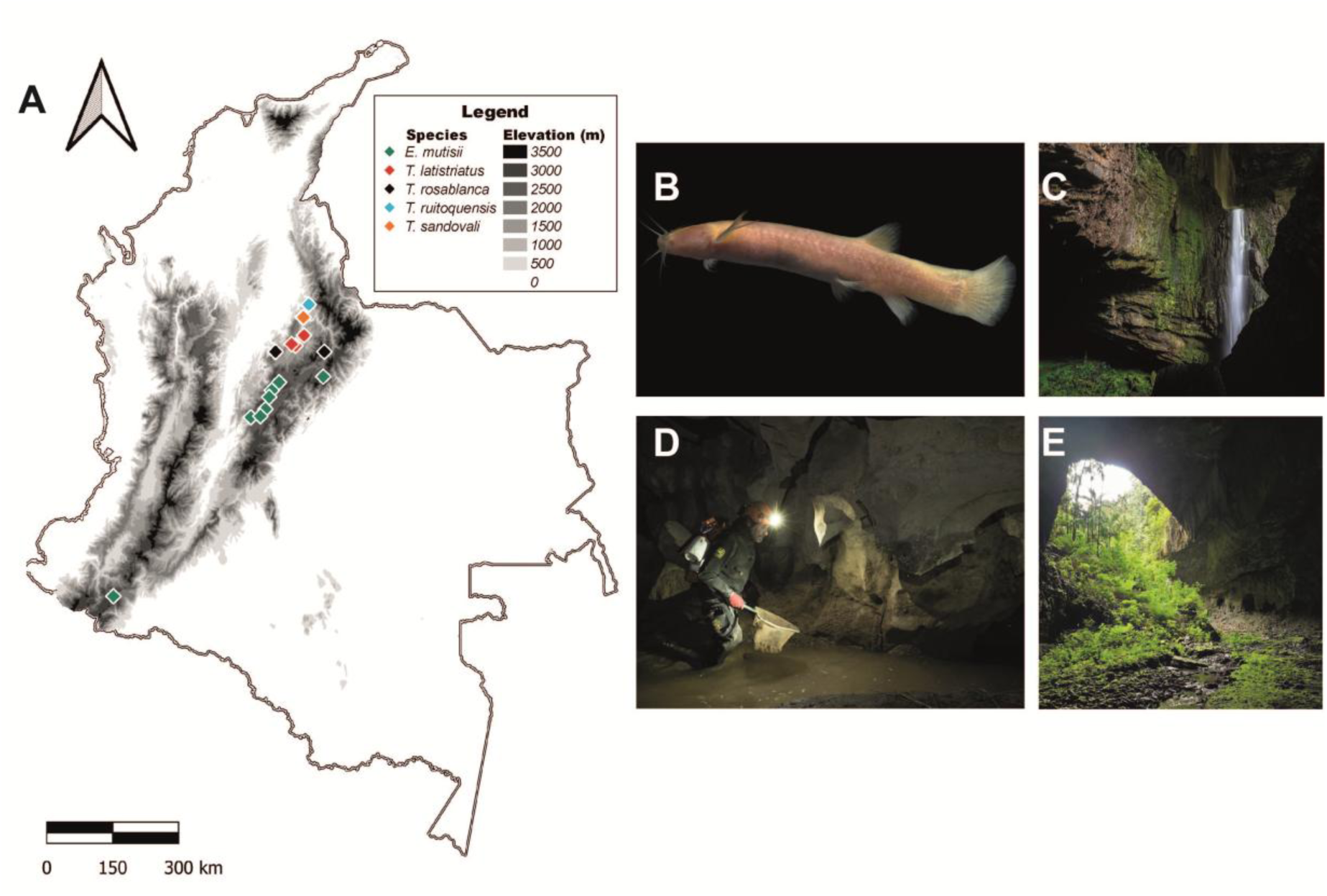
Geographic and environmental context of the *Trichomycterus* populations analyzed. (A) Collection localities of the examined specimens based on biological collection records, with sites coded according to species identity (see legend). (B) Live individual of *T. rosablanca* exhibiting troglomorphic traits. (C) Interior waterfall in El Peñón Cave, Santander, Colombia. (D) Fishing inside cave in Las Sardinas Cave, El Peñón, Santander, Colombia. (E) Cave entrance at La Concordia Cave, La Granja, Santander, Colombia. Photographs in panels B and C: Felipe Villegas Vélez, courtesy of the Instituto de Investigación de Recursos Biológicos Alexander von Humboldt. Photograph in panel D courtesy of Sofia Oggioni. Photograph in panel E courtesy of Carlos Lasso. All rights reserved.

### Body shape analysis

We used geometric morphometric analyses to investigate variation in body shape. A total of 236 specimens deposited in the freshwater fish collection of the Instituto de Investigación de Recursos Biológicos Alexander von Humboldt (IAvH-P) and the ichthyology collection at the Instituto de Ciencias Naturales at Universidad Nacional de Colombia were photographed in lateral view from the right side using a scale for reference.

We established 12 anatomically homologous landmarks following Colihueque *et al*. (2017) and Aguirre & Jiménez-Prado (2018), as shown in Table 2. To capture overall body outline variation, we complemented these landmarks with a dense sampling of the body contour using semi-landmarks (Luo, 2024; Monteiro, Guillermo, Rivera, & Di Beneditto, 2004). Using tpsDig2 software (Rohlf, 2017), we digitized manually a single continuous curve along the lateral outline of the body of each specimen, excluding fin extensions and caudal rays due to their variable preservation condition and positioning among specimens (Gunz & Mitteroecker, 2013; Mao et al., 2021). The curve was consistently traced in the same direction and initiated at landmark 1 (snout tip) for all individuals. Fifty equidistant points were then interpolated along this curve, providing a standardized representation of body outline shape. The combined landmark and outline semi-landmark configuration was used to describe global body shape variation (Fig. 2).

**Table 2.** Description of the anatomical landmarks used in the geometric morphometric analysis of *Trichomycterus* body shape. Each landmark corresponds to a distinct morphological reference point, and numbering matches the configuration presented in Fig. 2.

| Landmark | Description |
| --- | --- |
| 1 | Most anterior point of the snout |
| 2 | Most postero-ventral point of the operculum |
| 3 | Most antero-dorsal point of the operculum |
| 4 | Posterior margin of the head |
| 5 | Base of the pectoral fin |
| 6 | Anterior base of the dorsal fin |
| 7 | Anterior base of the anal fin |
| 8 | Posterior margin of the caudal peduncle in dorsal position |
| 9 | Posterior margin of the caudal peduncle in ventral position |
| 10 | Medial point of the caudal peduncle |
| 11 | Dorsal projection of LM5 to middle body, at the height of the lateral line |
| 12 | Ventral projection of LM7 to middle body, at the height of the lateral line |

**Figure 2.**
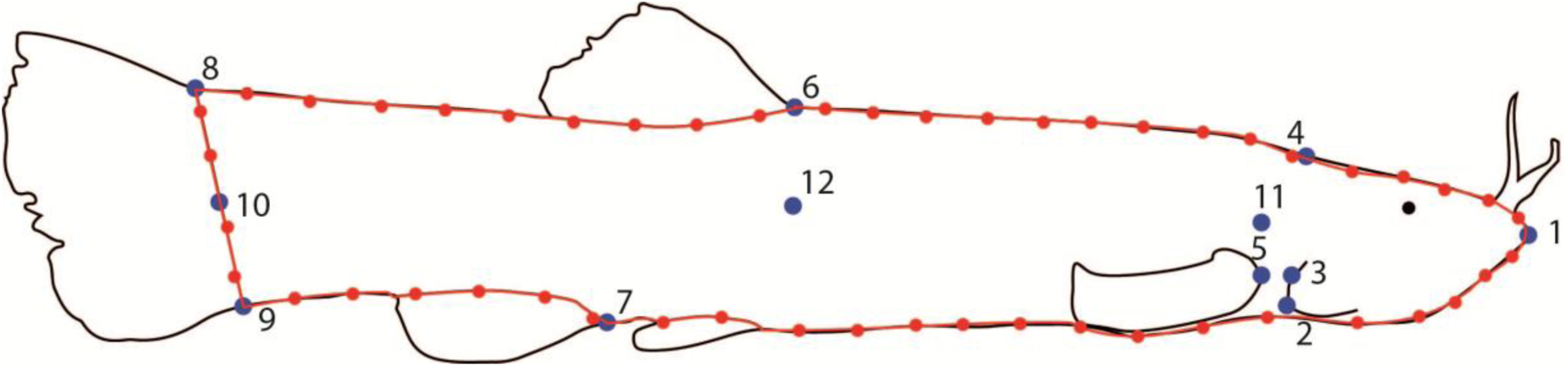
Landmark configuration used to quantify body shape in *Trichomycterus* catfishes. Image shows a lateral view of a specimen with 12 homologous landmarks (in blue), corresponding to anatomical reference points used in geometric morphometric analysis (see Table 2 for anatomical definitions). The red curve represents a body contour defined by 50 equidistant semilandmarks (in red) used to capture overall shape variation.

We digitized and converted landmarks into coordinates in a bidimensional plane using the tpsDig2 software (Rohlf, 2017). With the *‘Unbend specimens’* function in tpsUtil (Rohlf, 2004), we corrected the arching effect of the specimens (Valentin, Penin, Chanut, Sévigny, & Rohlf, 2008) using landmarks 1, 10, 11 and 12 to describe a quadratic curve used to remove the effect of curvature from all landmarks and generate a new set of coordinates for analyses.

The posture-corrected landmarks and outline coordinates obtained after the unbending procedure were aligned using a Generalized Procrustes Analysis (GPA), which scales all specimens to unit centroid size and removes differences due to translation and rotation based on a least-squares criterion, resulting in an optimal superimposition of configurations (Rohlf & Slice, 1990). This procedure isolates shape variation sensu stricto and allows direct comparison of overall body geometry among specimens.

To estimate measurement error, samples were digitized twice and evaluated through a Procrustes ANOVA, verifying that the mean squares associated with individuals were greater than those of the error (Arnqvist & Mårtensson, 1998). A Principal Component Analysis (PCA) was applied to the Procrustes-aligned coordinates as an exploratory ordination method to summarize the main axes of body shape variation and to visualize patterns of morphological similarity among specimens in relation to taxonomy and habitat of occurrence (Perazzo, Corrêa, Salzburger, & Gava, 2019).

We performed a multivariate regression of Procrustes-aligned landmark coordinates on the logarithm of centroid size, using a permutation test with 10,000 iterations to account for allometric shape variation (Drake & Klingenberg, 2008). The residuals of such regression, representing shape variation independent of size-related allometry, were retained for subsequent analyses and used to define a size-corrected morphospace through a PCA.

To examine habitat-and species-level patterns in body shape variation, we first analyzed individual scores along the first two principal components obtained from the PCA of size-corrected Procrustes coordinates, which accounted for 60% of the total variation to enable the visualization and interpretation of the main axes of morphological variation. In addition, to formally test for differences in overall body shape, we performed permutational multivariate analyses of variance (PERMANOVA; 10,000 permutations) using Euclidean distances on the matrix of size-corrected Procrustes residuals. Separate models were fitted to evaluate the effects of habitat (cave vs. surface) and species identity on body shape variation.

Finally, to determine whether cave and surface phenotypes are distinguishable based on their shape, we conducted a Discriminant Function Analysis (DFA) using habitat category as the *a priori* classification factor. The analysis was based on size-corrected shape variables and employed Mahalanobis distances, which account for the covariance structure among variables (Wang et al., 2025). Differences between group mean shapes were assessed using Hotelling’s T² statistic, which tests for multivariate separation between group centroids. In addition to the parametric T² test, significance was further evaluated using a permutation procedure (10,000 permutations), providing a non-parametric assessment of group differentiation. Geometric morphometric analyses, GPA, statistical and graphical analysis were performed using MorphoJ 1.07a (Klingenberg, 2011).

## 3 Results

The Procrustes ANOVA confirmed the reliability of landmark digitization, showing that the mean squares (MS) for individuals (4.885 × 10^−4^) greatly exceeded that associated with error (3 × 10^−5^). A significant allometric effect on body shape was detected (multivariate regression of Procrustes coordinates on centroid size, p < 0.001). Consequently, all subsequent analyses were conducted using size-corrected shape variables.

Permutational multivariate analyses of variance (PERMANOVA) based on Euclidean distances calculated from size-corrected Procrustes residuals revealed significant differences in overall body shape between habitats (F = 47.9, p < 0.001) and among species (F = 14.07, p < 0.001). These results indicate that body shape variation is structured both by habitat and by species identity.

The principal component analysis of shape variation revealed consistent patterns of association with habitat and species identity (Fig. 3). The first two principal components accounted for 59.62% of total shape variation (PC1 = 34.66%, PC2 = 24.96%). Along PC1, specimens primarily varied in body elongation and overall fusiformity, whereas PC2 captured variation related to mid-body depth, caudal peduncle width, and the anteroposterior position of the pectoral fins. In the morphospace defined by these axes, cave-dwelling specimens tended to occupy regions characterized by more fusiform bodies with shallower body depth and wider caudal peduncles, whereas surface-dwelling specimens were generally associated with morphologies with deeper, more robust bodies and narrower caudal peduncles (Fig. 3).

**Figure 3.**
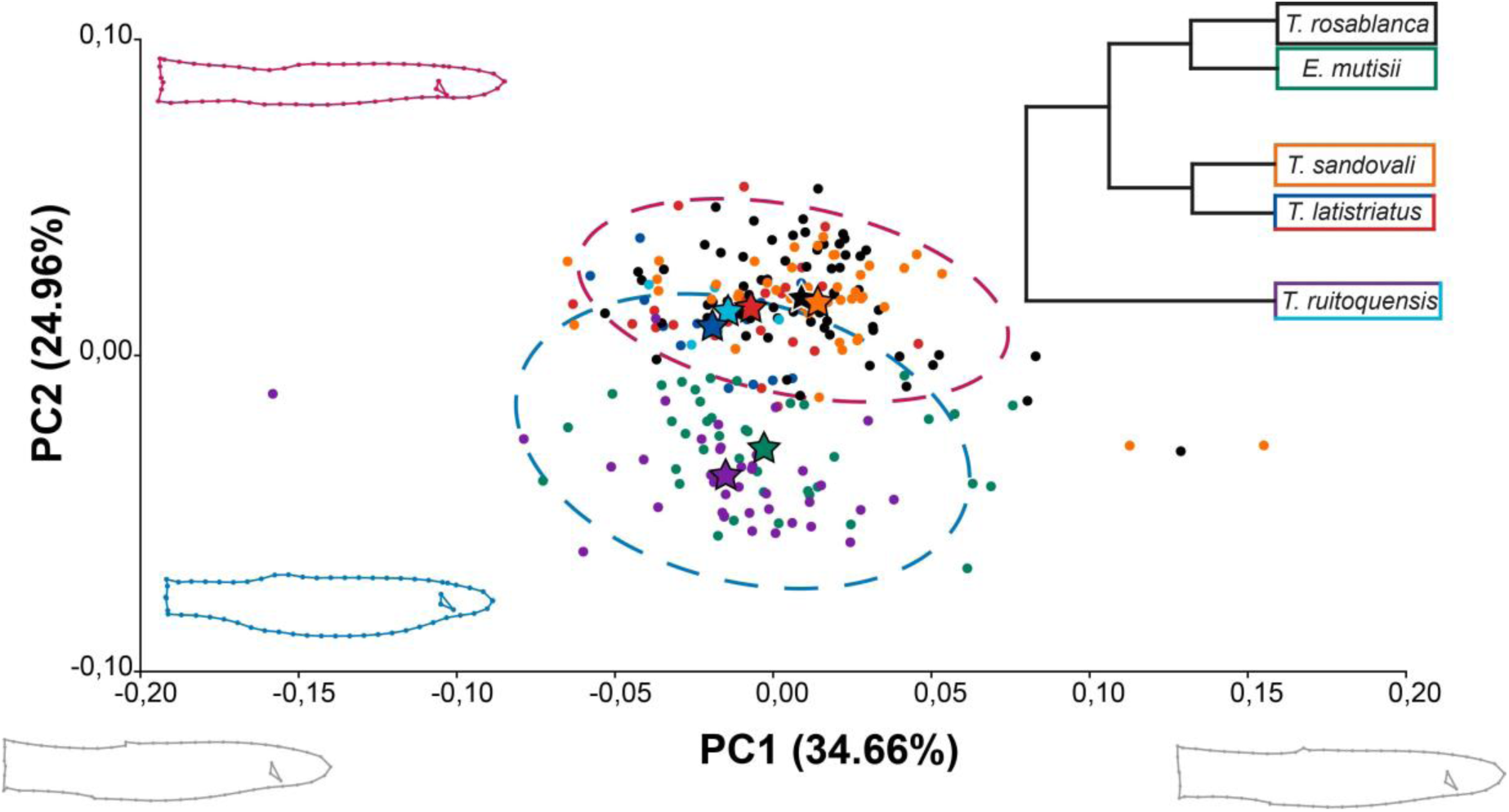
Cave and surface *Trichomycterus* species occupy distinct regions of morphospace, revealing consistent shape differences across habitats. Scatterplot of PC1 vs. PC2 based on size-corrected body shape variables; each point represents an individual, colored by species. Dashed ellipses represent 90% confidence intervals for habitat groups: cave-dwelling species (magenta) and surface-dwelling species (blue). Stars indicate species centroids (mean position of each species in morphospace). Silhouettes depict mean body shapes at the extremes of PC2, illustrating habitat-associated divergence, while variation along PC1 is minor. The internal triangle denotes the operculum–pectoral fin relationship (see Table 2). The simplified phylogeny (after Flórez et al. 2021) shows species relationships. Box colors indicate species identity. In two species with both habitat types (*T. latistriatus* and *T. ruitoquensis*), box colors distinguish populations: cave (red for *T. latistriatus*, light blue for *T. ruitoquensis*) and surface (dark blue for *T. latistriatus*, purple for *T. ruitoquensis*). Remaining species are represented by single colors: *E. mutisii* (green), *T. rosablanca* (black), and *T. sandovali* (orange).

To evaluate whether the patterns observed in the PCA ordination were statistically supported individually, habitat-related differences were further examined using the distribution of individual PC scores (Fig. 4). Comparisons between cave and surface specimens revealed a statistically significant difference along PC1 (p < 0.001), although the distributions showed substantial overlap between habitats, indicating that this axis captures limited biologically meaningful differentiation. In contrast, PC2 exhibited clearer separation between cave and surface specimens (p < 0.001), supporting its relevance as the primary axis of habitat-associated shape variation. When PC scores were examined by species, cave populations (e.g., *Trichomycterus rosablanca*, *T. sandovali* and *T. ruitoquensis* from caves) exhibited broadly overlapping distributions consistent with their shared habitat, while surface populations (*Eremophilus mutisii* and *T. ruitoquensis* from surface) displayed similar patterns distinct from those of cave taxa. *Trichomycterus latistriatus* showed a broad and overlapping distribution across axes, with no clear differentiation relative to either cave or surface groups. Although outliers were more frequent in surface habitats and in some species (e.g., *T. latistriatus*), their exclusion did not affect the results, indicating that the observed patterns are robust to these observations.

**Figure 4.**
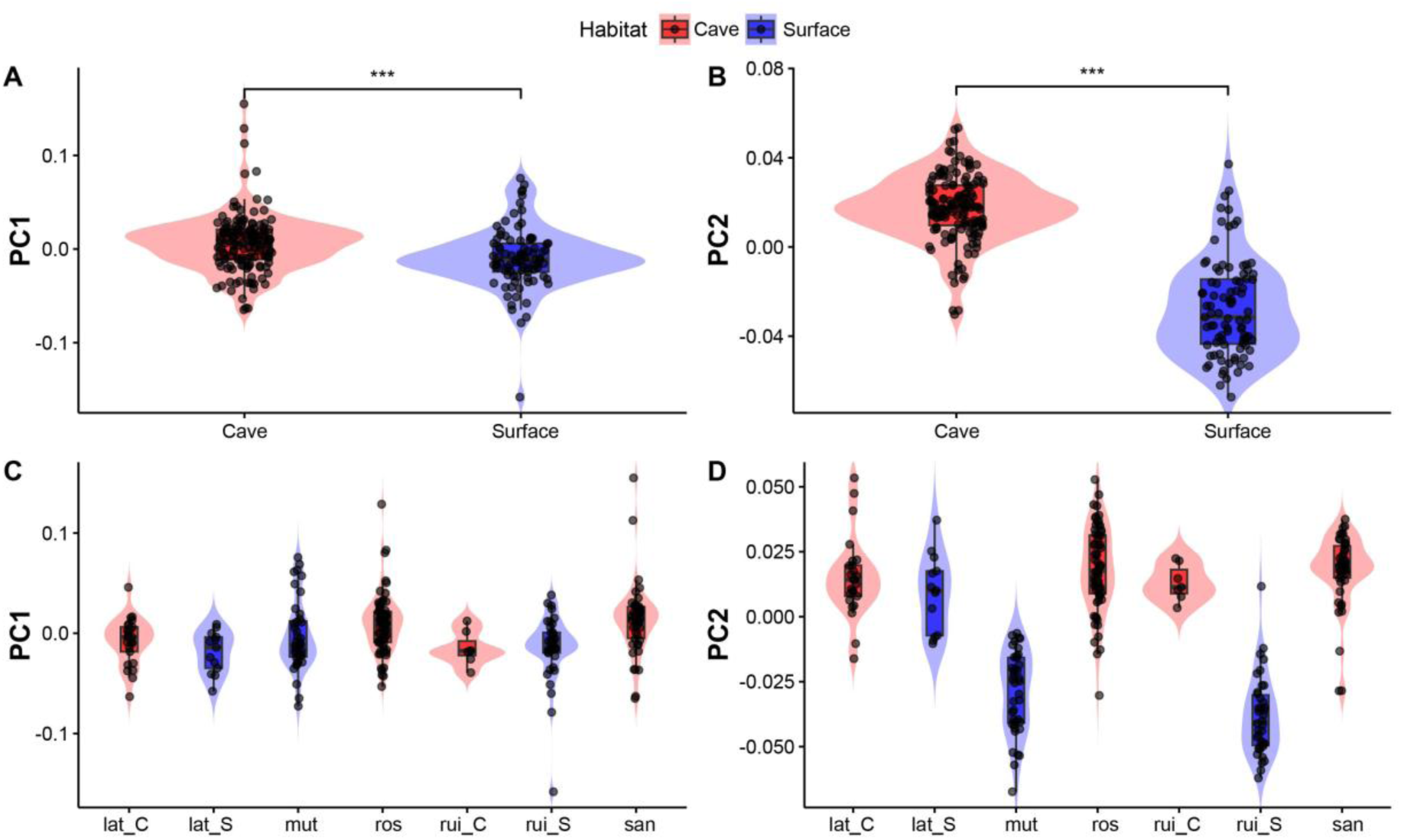
Cave- and surface-dwelling *Trichomycterus* differ in body shape along the first two principal components. Panels A–B show habitat-level comparisons of PC1 and PC2 scores, respectively, with significant differences between cave (red) and surface (blue) groups (p < 0.001). Panels C–D show species-level variation in PC1 and PC2 scores across habitats, with violin plots, boxplots, and individual data points for each group, revealing consistent habitat-associated patterns within species. Colors indicate habitat (red = cave, blue = surface). In *T. latistriatus* and *T. ruitoquensis*, suffixes denote habitat-specific populations (_C = cave, _S = surface). Species abbreviations: mut = *E. mutisii*, lat = *T. latistriatus*, ros = *T. rosablanca*, rut = *T. ruitoquensis*, san = *T. sandovali*.

Finally, discriminant function analysis based on size-corrected shape variables showed that 100% of cave specimens and 98% of surface specimens were correctly classified according to the habitat where the specimens were captured. Differences between habitat group centroids were statistically significant (Hotelling’s T² test, *p* < 0.001), indicating that overall body shape allows discrimination between cave and surface phenotypes.

## 4 Discussion

Our study provides new evidence of morphological differentiation between cave-dwelling and surface-dwelling trichomycterid catfish species in Colombia that extends beyond the pigmentation and eye development variation emphasized in earlier work. While previous studies on cave fishes have largely focused on regressive traits such as eye reduction, depigmentation, and craniofacial modifications (e.g., Holtz & Albertson, 2024; Powers et al., 2017; Soares & Niemiller, 2020), our results reveal consistent habitat-associated differences in overall body shape across multiple species and lineages, highlighting an underexplored dimension of phenotypic variation in Trichomycteridae. Differences are not restricted to interspecific comparisons but also emerge within species that occupy both cave and surface environments, suggesting that body shape variation is consistently associated with habitat conditions, although with varying strength among lineages. This pattern is indicative of repeated morphological differentiation linked to habitat use, rather than isolated species-specific divergence.

The observed variation in body shape among *Trichomycterus* species is consistent with habitat-associated functional differentiation, potentially linked to contrasting hydrodynamic conditions in cave and surface environments. Although trichomycterid catfishes are generally described as elongated and narrow-bodied (Adriaens, Baskin, & Coppens, 2010), our analyses revealed recurrent and directionally consistent differences in body configuration between habitats, particularly along axes associated with elongation, body depth and caudal peduncle width (Figs. 3-4). These differences, while subtle at the level of individual specimens, are clearly expressed in shape configurations and correspond to significant divergence, especially along PC2, which captures the main axis of habitat-associated shape variation. Cave-dwelling species from independent lineages exhibited similar fusiform morphologies, anteriorly positioned pectoral fins, and wider caudal peduncles, whereas surface-dwelling species displayed deeper, more compressed bodies with posteromedially positioned fins (Fig. 3). However, the strength and expression of this pattern vary among species: while some taxa exhibit clear separation between habitats, others (e.g., *T. latistriatus*) retain similar morphologies across different environments, suggesting that habitat-associated shape divergence may be modulated by lineage-specific constraints or evolutionary history. These patterns indicate a general association between habitat type and body shape, which could be indicative of habitat-mediated phenotypic convergence, which was not formally quantified in this study.

At first glance, the fusiform body shapes observed in cave-dwelling species appear to contradict classical expectations that structurally complex environments favor deeper, more compressed morphologies associated with enhanced maneuverability (Langerhans, 2008; Langerhans & Reznick, 2010; Webb, 1984). Rather than reflecting a contradiction, this pattern may arise from differences in the ecological conditions experienced by fishes in cave and surface systems across multiple dimensions, including structural complexity, hydrodynamics, resource availability, and biotic interactions. While many surface benthic habitats include dense vegetation, heterogeneous substrates, and variable flow regimes, subterranean aquatic systems are often characterized by reduced primary productivity and the absence of macrophytes (Lasso et al., 2019), although they may still present considerable structural complexity. These differences suggest that the functional demands imposed by cave environments may not directly parallel those of structurally complex surface habitats and could instead favor alternative locomotor strategies. In such context, the fusiform morphology observed in cave-dwelling species is consistent with expectations from hydrodynamic theory (Langerhans & Reznick, 2010) and likely reflect the combined influence of multiple ecological factors rather than a single dominant driver. Also, we note that the fusiform shapes in cave populations of *Trichomycterus* we documented are not universal across troglomorphic species in the genus occupying such environments in Colombia (e.g. *T. spectrum* from the Río Ranchería basin; DoNascimiento & Prada-Pedreros, 2020).

Our study does not directly measure hydrodynamic conditions or swimming performance and therefore cannot establish a direct functional link between body shape and locomotor efficiency. However, the observed morphological patterns are best understood as emerging from the interaction of multiple ecological factors, including hydrodynamics, habitat structure and resource availability. The cave–surface contrast examined here should thus be interpreted as a broad ecological gradient encompassing diverse and potentially heterogeneous conditions across systems. Within this framework, the repeated occurrence of similar body shapes in cave-dwelling lineages is consistent with patterns of convergent evolution widely documented in subterranean fishes, even if the specific selective pressures driving this convergence remain unresolved. Due to the repeated association between habitat and shape suggest that functional constraints are linked to habitat structure, the functional interpretations proposed here should be viewed as testable hypotheses rather than direct evidence of adaptation (Losos, 2011; Moen, 2019; Walker, 2007).

Future research should explicitly test these hypotheses by integrating biomechanical and ecological data into an evolutionary framework to further investigate the functional and evolutionary mechanisms underlying the patterns documented here. Experimental approaches such as flow-tunnel assays, maneuverability tests in structurally complex environments, common-garden experiments and trait-specific responses to life in caves could help determine whether the body shape differences documented here translate into performance trade-offs and whether phenotypic plasticity contributes to the observed patterns (Holzman, Perkol-Finkel, & Zilman, 2014; Sampaio, Rufino, Pompeu, de Andrade e Santos, & Ferreira, 2020; Yoffe et al., 2020). Such approaches would allow disentangling adaptive differentiation from plastic or constraint-driven responses, particularly in systems where direct ecological measurements are challenging.

Subterranean habitats are known to impose strong and recurrent selective pressures that can lead to both regressive and constructive traits, often resulting in similar phenotypes across independent lineages (Losos, 2011; Moldovan, Kováč, & Halse, 2019; Soares & Niemiller, 2020; Trontelj, Blejec, & Fiser, 2012). While regressive traits in cave fishes have been extensively documented, our results indicate that body shape also exhibits consistent, habitat-associated variation across species, suggesting that locomotor-related morphology may represent an additional, yet underexplored, axis of phenotypic differentiation in subterranean systems. In particular, the repeated association between fusiform body shapes and cave environments observed here is consistent with broader patterns of morphological convergence, although the strength and consistency of this pattern vary among lineages, indicating that lineage-specific constraints and evolutionary histories may play an important role in shaping morphological outcomes. Our results therefore extend previous work by integrating body shape into the suite of traits associated with subterranean life, complementing previously described cranial, vertebral and sensory modifications (Chen et al., 2022; Mao et al., 2021; Powers et al., 2017; W. Reyes & Aguirre, 2019).

*Trichomycterus latistriatus* occupies an intermediate position in morphospace, consistent with its occurrence in both cave and surface environments. However, individuals from different habitats do not fully separate in morphospace, and instead cluster around a shared morphological configuration (Fig. 3), suggesting that body shape in this taxon may be less responsive to local habitat conditions than in other species analyzed. This pattern contrasts with that observed in *T. ruitoquensis*, where individuals from cave and surface environments show greater differentiation, indicating that the relationship between habitat and body shape is not uniform across taxa. Taken together, these results suggest that habitat-associated morphological variation may be modulated by lineage-specific factors, such as evolutionary history or functional constraints, rather than reflecting a simple or universal response to environmental conditions.

Rather than representing a strictly “intermediate” morphology, the body shape of *T. latistriatus* may reflect a more constrained configuration, potentially limiting its divergence across environments. This interpretation is consistent with the broader pattern observed here, where convergence toward cave-associated morphologies is evident but not ubiquitous across all lineages. Given ongoing taxonomic and phylogenetic revisions within Andean *Trichomycterus* (Arratia, 1990; Fernandez et al., 2021), future work should clarify whether the observed morphological variation reflects ecological flexibility within a single lineage or cryptic diversity associated with habitat specialization.

## 5 Conclusions

Overall, our findings document consistent, habitat-associated patterns of body shape variation in Colombian trichomycterid catfishes, with cave-dwelling species exhibiting more fusiform body shapes and surface-dwelling species exhibiting deeper bodies. Although we do not directly test for convergent evolution or functional performance, the recurrence of similar morphologies among cave-dwelling species from independent lineages is consistent with repeated, though not uniform, phenotypic responses to shared ecological constraints. These results highlight the value of integrating geometric morphometrics with ecological and biomechanical approaches to better understand the evolutionary processes shaping phenotypic diversity in subterranean and surface aquatic systems.

## Acknowledgments

We thank the Facultad de Ciencias at Universidad de los Andes, which funded this project through a ‘Proyecto Semilla’ research grant to Nelson Falcón Espitia. We would like to thank the curators of the collections visited (IAvH-P and ICN-P) and Laura Pabón for their help and technical assistance during this project. The authors truly appreciate the valuable input from Jorge Molina, Juan Camilo Ríos, Andrés Link, Hernán López-Hernández, and Carlos DoNascimiento during the development of this project. We also thank Felipe Villegas, Carlos Lasso (IAvH) and Sofia Oggioni for the photographs of *Trichomycterus rosablanca* and cave habitats. Finally, we thank the two anonymous reviewers for their comments which improved the quality of this manuscript.

## Author contribution statement

The study was conceptualised by NFE and CDC. NFE developed the methodology, conducted the investigation, and performed the data analysis. Data interpretation was carried out by NFE and CDC. NFE prepared the figures and tables. The manuscript was written by NFE and CDC. All authors critically revised the manuscript and approved the final version for submission.

## Data Availability Statement

Data is available upon request.

